# Computed tomography reveals hip dysplasia in the extinct Pleistocene saber-tooth cat *Smilodon*

**DOI:** 10.1101/2020.01.07.897348

**Authors:** Mairin A. Balisi, Abhinav K. Sharma, Carrie M. Howard, Christopher A. Shaw, Robert Klapper, Emily L. Lindsey

## Abstract

Reconstructing the behavior of extinct species is challenging, particularly for those with no living analogues. However, damage preserved as paleopathologies on bone can record how an animal moved in life, potentially reflecting behavioral patterns. Here, we assess hypothesized etiologies of pathology in a pelvis and associated right femur of a *Smilodon fatalis* saber-toothed cat, one of the best-studied species from the Pleistocene-age Rancho La Brea asphalt seeps, California, USA, using visualization by computed tomography (CT). The pelvis exhibits massive destruction of the right hip socket that was interpreted, for nearly a century, to have developed from trauma and infection. CT imaging reveals instead that the pathological distortions characterize chronic remodeling that began at birth and led to degeneration of the joint over the animal’s life. These results suggest that this individual suffered from hip dysplasia, a congenital condition common in domestic dogs and cats. This individual reached adulthood but could not have hunted properly nor defended territory on its own, likely relying on a social group for feeding and protection. While extant social felids are rare, these fossils and others with similar pathologies are consistent with a spectrum of social strategies in *Smilodon* supported by a predominance of previous studies.

## Introduction

The saber-toothed cat *Smilodon fatalis* is one of the best-studied apex predators from the late Pleistocene, if not across the entire history of fossil mammals^1^. Much of our knowledge about this species comes from the Rancho La Brea (RLB) asphalt seeps in Los Angeles, California, United States, which have preserved thousands of individuals of *Smilodon* from at least 50,000 years ago until their extinction approximately 11,000 years ago^1^. The seeps primarily functioned as a carnivore trap: a large herbivore mired in the asphalt inadvertently would attract large carnivores and scavengers, which themselves would become entrapped in great numbers^2^. Studies of *Smilodon* at RLB have enabled reconstruction of its feeding behavior as an ambush predator specializing on herbivorous megafauna, inferred using independent approaches ranging from comparative morphology (e.g., ^3^) to stable isotopes (e.g., ^4^). As well, the abundant specimens include numerous examples of bone lesions, or pathologies^5^: a phenomenon that tends to be rare at more typical fossil sites (e.g., fluvial deposits, bone beds, ashfalls, caves) but captured by RLB’s large sample sizes.

Because bone remodels throughout an animal’s life in response to stress, strain, and injury^6^, paleopathologies can preserve a record of realized behavior and supplement the picture of potential behavior presented by skeletal morphology. Differences in the distribution of pathologies throughout the skeleton, for example, distinguish *Smilodon* from a contemporaneous predator, the dire wolf *Aenocyon* (“*Canis*”) *dirus*, reflecting interspecific differences in hunting behavior and prey preference corroborated by independently gathered data^4^. While injuries in the dire wolf tend to be concentrated around its distal limbs, supporting the hypothesis that it was a pursuit predator, *Smilodon*’s injuries cluster around its midline, supporting inferences that it ambushed and grappled with prey^5^. As the aggregate result of how an animal moved over the course of its life, pathologies present a relatively direct record of the animal’s interactions with its predators, prey, environment, and even conspecifics, potentially including intraspecific interactions like social behavior.

The current study centers on a *Smilodon* specimen (LACMHC 131) that has been described as “the most strikingly pathological object in the collection of Rancho La Brea fossils”^7^. The specimen is a right innominate bone exhibiting massive distortion and destruction of the hip socket (Fig. 1). An analysis in 1930 by Moodie^7^, restricted to an inspection of the gross morphology, proposed that this pathology had resulted from infection following violent trauma, possibly during an encounter with a conspecific, which also led to dislocation of the femur from the hip. Moodie found no pathological bones potentially associated with the injured innominate; because of disarticulation by flowing asphalt over thousands of years, associated elements are rarely encountered at RLB. However, more than half a century later, Shermis^8^ described a pathological femur (LACMHC 6963; Fig. 2) associated with another pathological pelvis. Later, this femur was determined to be associated instead with the specimen described by Moodie^9^, enabling examination of the effects of a single event on associated skeletal elements.

**Figure 1.**
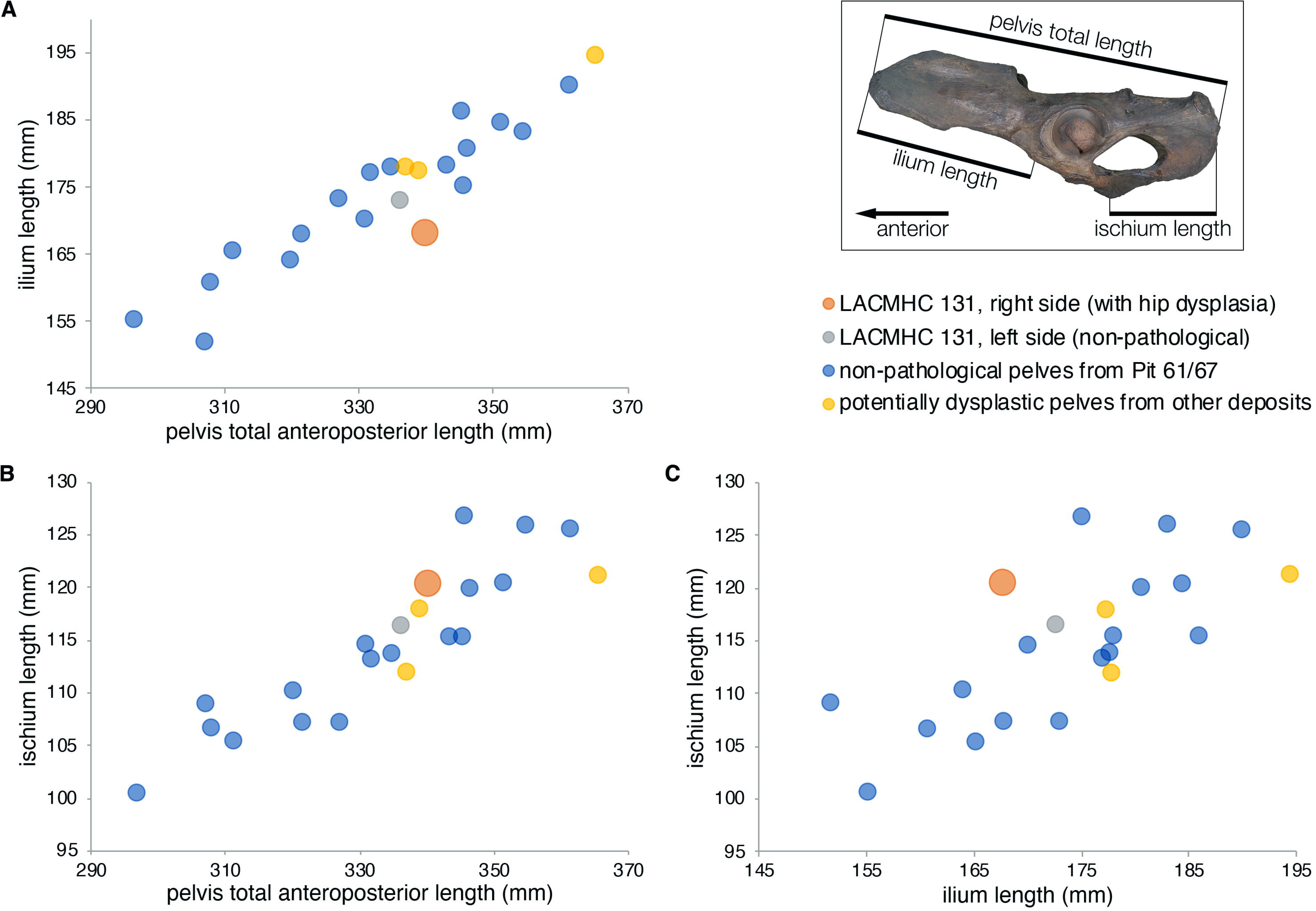
Photographs of LACMHC 131, a pathological pelvis belonging to *Smilodon fatalis*. **(A)** Right lateral view showing destruction of the acetabulum; anterodorsal end to the right. **(B)** Left lateral view showing the acetabulum intact but with exostoses around the anterodorsal rim; anterodorsal end to the left. **(C)** Dorsal and **(D)** ventral views showing asymmetry in the pelvis; anterior end to the right.

**Figure 2.**
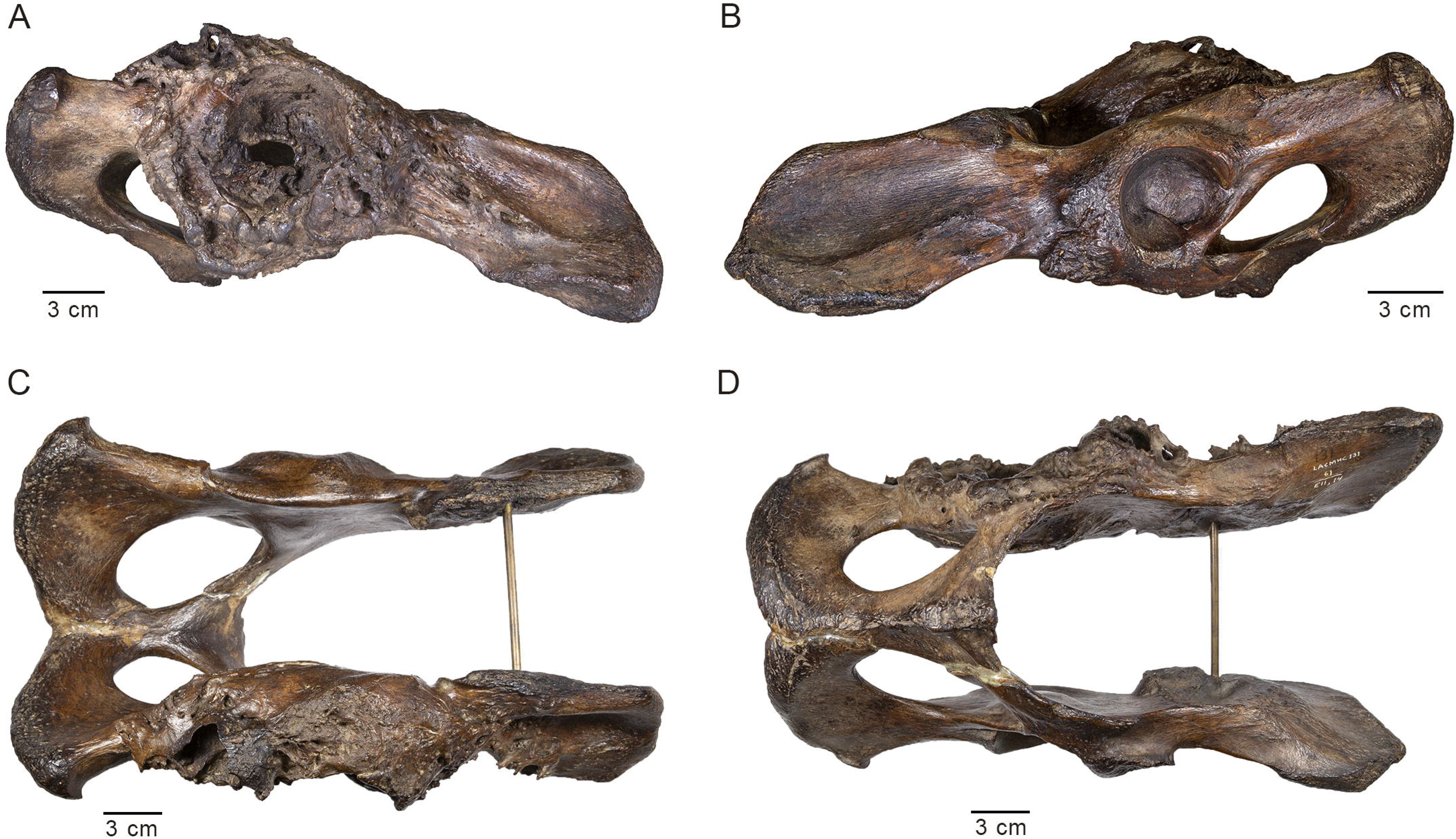
Photographs of LACMHC 6963, a pathological right femur belonging to *Smilodon fatalis*. **(A)** Anterior and **(B)** posterior views of full femur, excluding the distal epiphysis broken after death; proximal end on the left. **(C)** Anterior and **(D)** posterior close-up views of the proximal end. **(E)** Dorsal close-up view of the femoral head, greater trochanter, and lesser trochanter in lower center background. **(F)** Lateral close-up view of the greater trochanter and lesser trochanter (lower center), which is enlarged into a round knob. The upper scale bar refers to A and B and the lower scale bar refers to C, D, E, and F.

In the present study, we supplement gross morphology and analyze LACMHC 131 and 6963 using computed tomography to observe the internal bone structure of these elements. We evaluate the historical inference that this pathology was the result of trauma and assess different etiologies: traumatic arthritis, infective arthritis, or degenerative arthritis. We present basic morphometric data, quantitatively compare these pathological bones to other pathological and non-pathological *Smilodon* elements at RLB, and examine these specimens in the context of similar pathologies in living cats. Finally, we explore the implications of the diagnosis on reconstructions of social behavior in *Smilodon* and the potential contribution of paleopathology to a growing interdisciplinary body of literature supporting sociality in this extinct predator.

## Results

### Description

LACMHC 131 is an intact pelvis with all sutures completely fused (Fig. 1; Supplementary File S1; Supplementary Video S1). The distal end of LACMHC 6963, the femur, is broken and missing (Fig. 2; Supplementary File S1), precluding verification of distal epiphyseal fusion. However, the proximal epiphyses are fused to the shaft.

Imaging shows no signs of callus, or bone regeneration and healing, typically seen following fracture. Rather, osteophytes likely signal bone remodeling secondary to malformation of the joint. The changes in the right acetabulum and right femur are consistent with those expected from repetitive subluxation and subsequent necrosis. The right acetabulum is shallow and elliptical-shaped as opposed to concentric-shaped. A hole in the bone, likely the result of postmortem asphaltic wear given its sharp edges, marks the thin medial wall of the acetabulum, which is lined otherwise with exostoses. The left acetabulum appears non-pathological; however, the ilium anterodorsal to the socket—origin of the quadriceps femoris muscles—is rugose (Fig. 1; Supplementary File S1; Supplementary Video S1), contrasting with typical *Smilodon* pelvic specimens (e.g., Supplementary Fig. S1). The head of the pathological right femur is flattened and laden with anatomical distortions; as well, the lesser trochanter is enlarged into a round knob (Fig. 2; Supplementary File S1). In contrast, a non-pathological femur bears a round head that is appropriately developed (Supplementary Fig. S2). This head would fit snugly into a concentric-shaped socket such as the left acetabulum of the pathological pelvis or either acetabulum of a non-pathological pelvis (e.g., Supplementary Fig. S1), allowing for a normal axis of rotation and movement.

The four cardinal findings of arthritis on imaging are bony sclerosis, osteophytes, joint space narrowing, and subchondral cysts. The CT images of the pathological specimens reveal evidence of such degenerative changes and an absence of fractures from traumatic impact (Figs. 3, 4; Supplementary Video S2; Supplementary Data S1-S3). The images demonstrate sclerosis and osteophytes in the right acetabulum (Fig. 3), which are changes consistent with degenerative arthritis. Profuse remodeling with osteophyte formation also marks the right femoral head (Fig. 4), likely in response to the degenerative process from repeated subluxation and dysplasia. This contrasts with the relatively smooth, evenly dense cortical bone of the non-pathological femur (Fig. 5). Altogether, these findings indicate dysplasia of the hip as the pathological cause.

**Figure 3.**
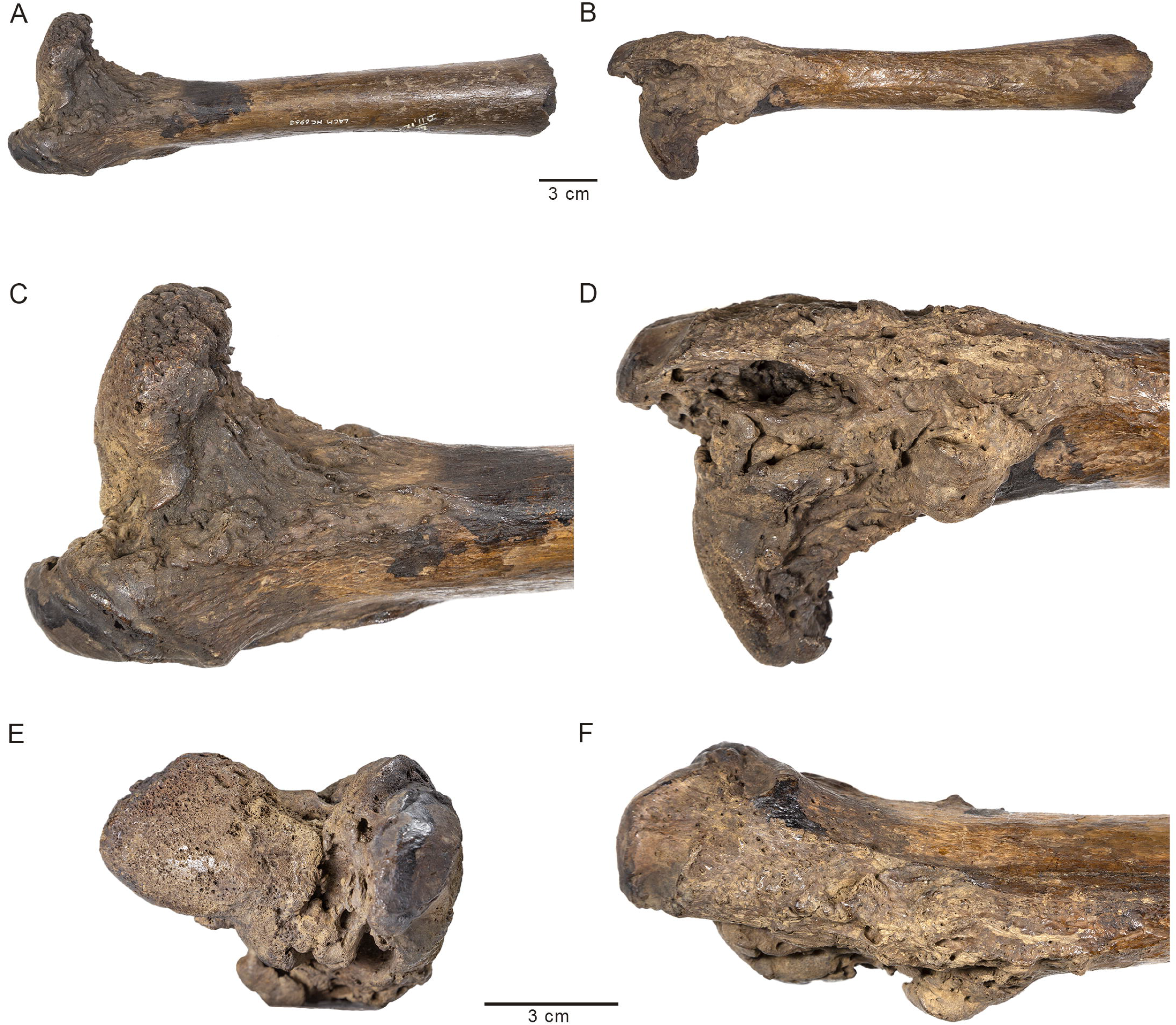
Three-dimensional model of the pathological *Smilodon* pelvis and acetabulum (LACMHC 131) reconstructed from CT images. Horizontal numbered lines in (A) mark anatomical location of numbered CT slices displayed in (E). **(A)** Ventral view of pelvis, anterior toward top of image; **(B)** rotated 45 degrees from ventral about the anteroposterior axis; **(C)** rotated 90 degrees from ventral about the anteroposterior axis; **(D)** rotated 135 degrees from ventral about the anteroposterior axis. **(E)** 0.63mm axial CT images of the pathological pelvis demonstrating a dysplastic right hip socket, corresponding with the three-dimensional model, posterior view.

**Figure 4.**
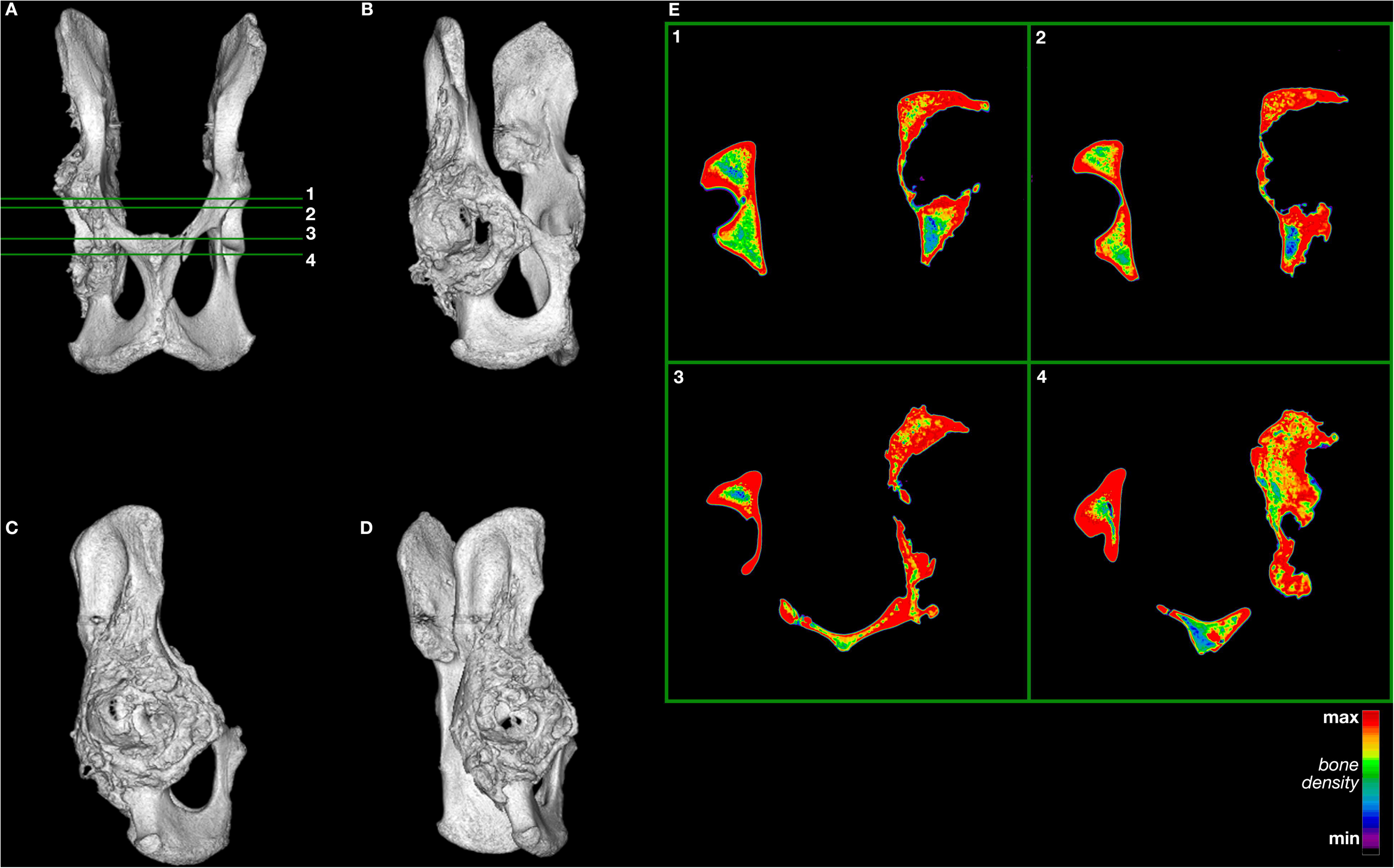
Three-dimensional model of the pathological *Smilodon* right femur reconstructed from CT images. Horizontal numbered lines in (A) mark anatomical location of numbered CT slices displayed in (E). **(A)** Anterior view of right femur; **(B)** rotated 90 degrees from anterior about the vertical axis (lateral view); **(C)** rotated 180 degrees from anterior about the vertical axis (posterior view); **(D)** rotated 270 degrees from anterior about the vertical axis, displaying the femoral head and neck (medial view). **(E)** 0.63mm axial CT images of the pathological right femur demonstrating morphological distortion at the femoral head and neck, corresponding with the three-dimensional model. Each inset image has been rotated so that top corresponds to anterior, right corresponds to anatomical right, bottom corresponds to posterior, and left corresponds to anatomical left sides.

**Figure 5.**
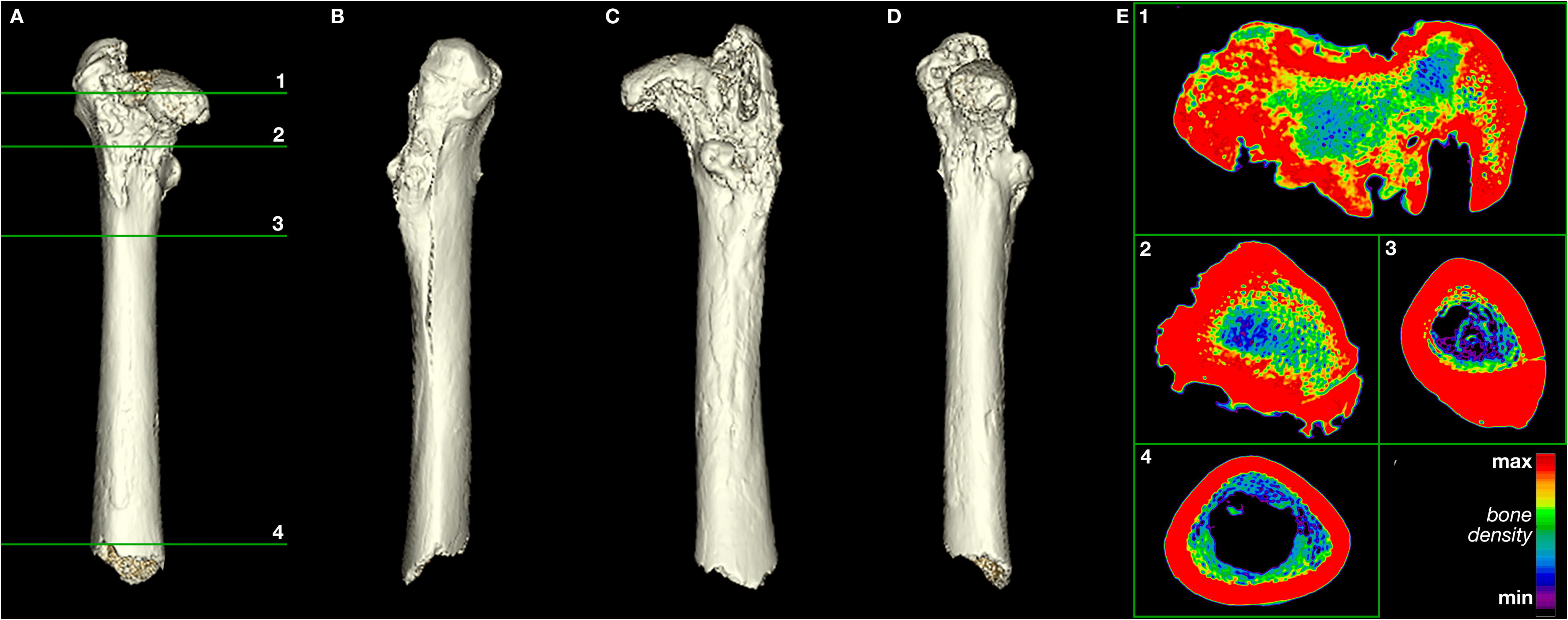
Three-dimensional model of a normal *Smilodon* right femur (LACMHC K-2584) reconstructed from CT images. Horizontal numbered lines in (A) mark anatomical location of numbered CT slices displayed in (E). **(A)** Anterior view of right femur; **(B)** rotated 90 degrees from anterior about the vertical axis (lateral view); **(C)** rotated 180 degrees from anterior about the vertical axis (posterior view); **(D)** rotated 270 degrees from anterior about the vertical axis, displaying the femoral head and neck (medial view). **(E)** 1.25mm axial CT images of the normal right femur corresponding with the three-dimensional model. Each inset image has been flipped and rotated so that top corresponds to anterior, right corresponds to anatomical right, bottom corresponds to posterior, and left corresponds to anatomical left sides.

### Body-size estimation

Basic morphometric data for the two pathological specimens are presented in Table 1. The two specimens lie well within the size range of adult *Smilodon* pelves (Fig. 6) and femora (Fig. 7). These size comparisons as well as epiphyseal fusion suggest that the individual represented by these two specimens was an adult.

**Table 1.**
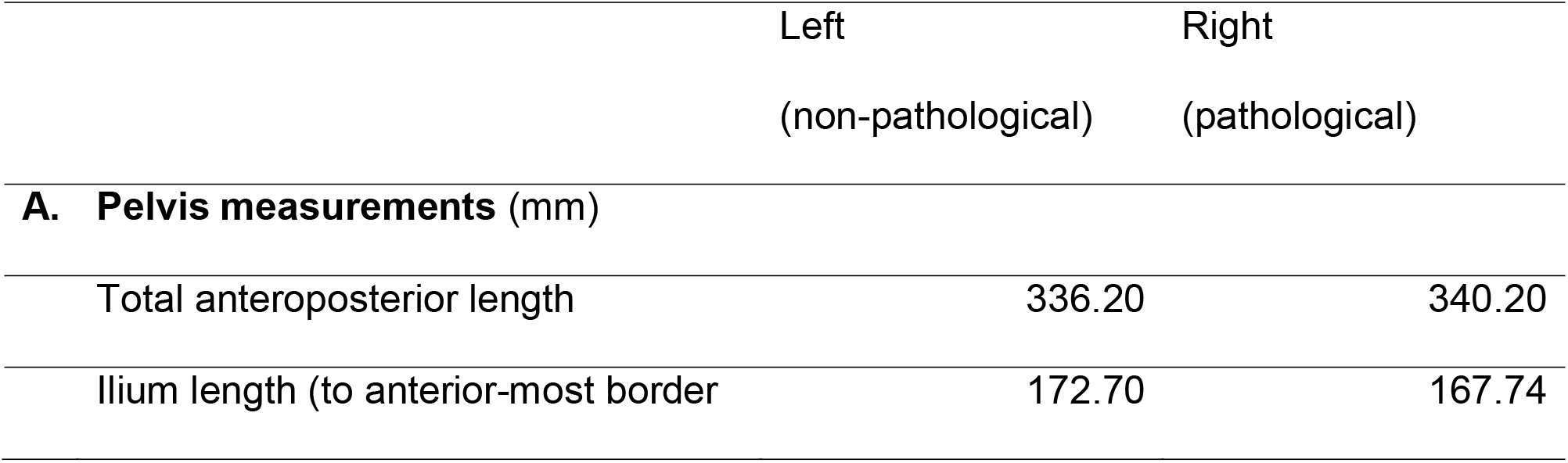
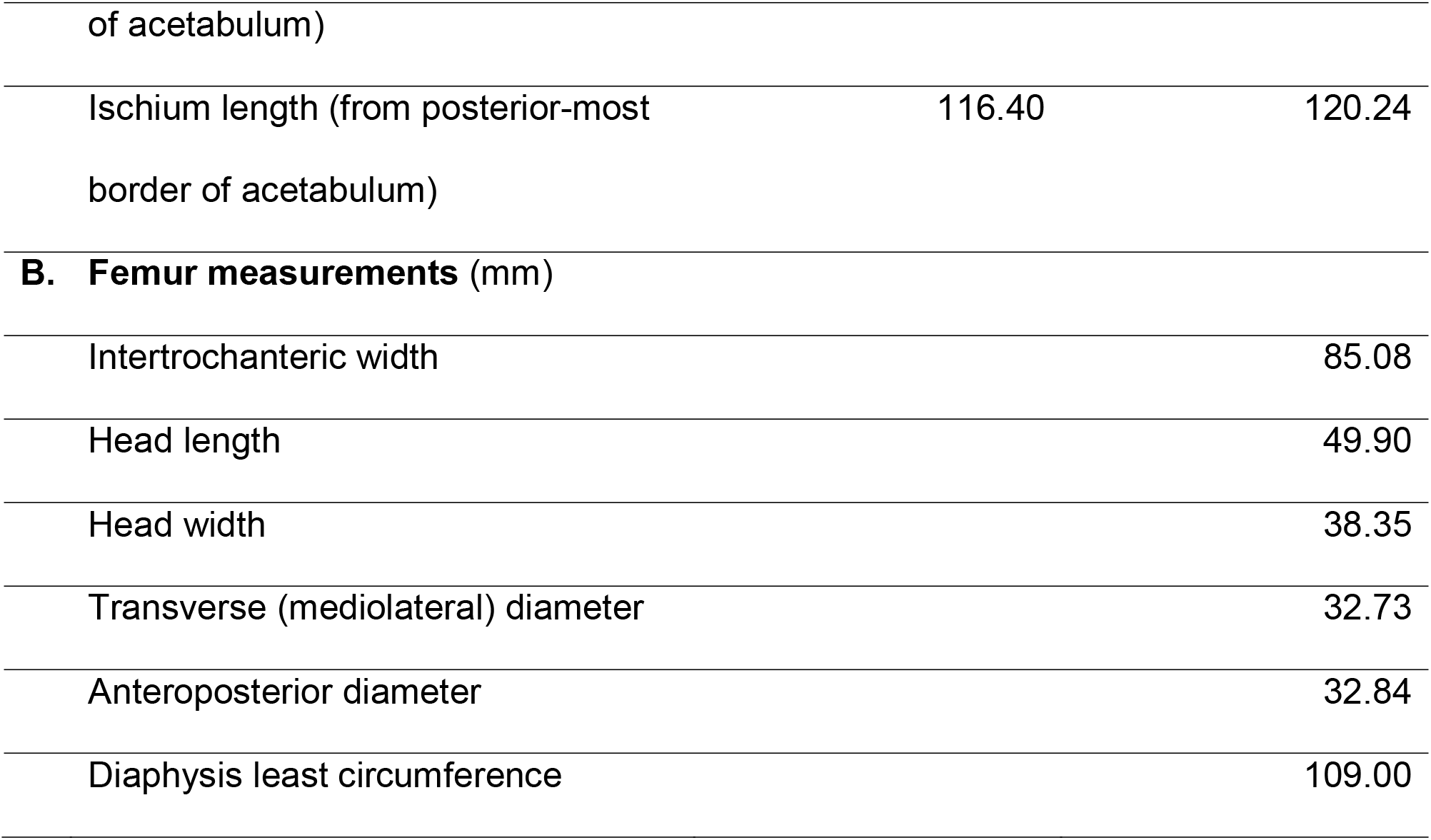
Basic morphometric data for the two focal pathological *Smilodon fatalis* specimens: (A) pelvis, LACMHC 131; (B) right femur, LACMHC 6963. Morphometric data for comparative specimens are in Supplementary Data S4 and S5 and plotted in Figures 6 and 7.

**Figure 6.**
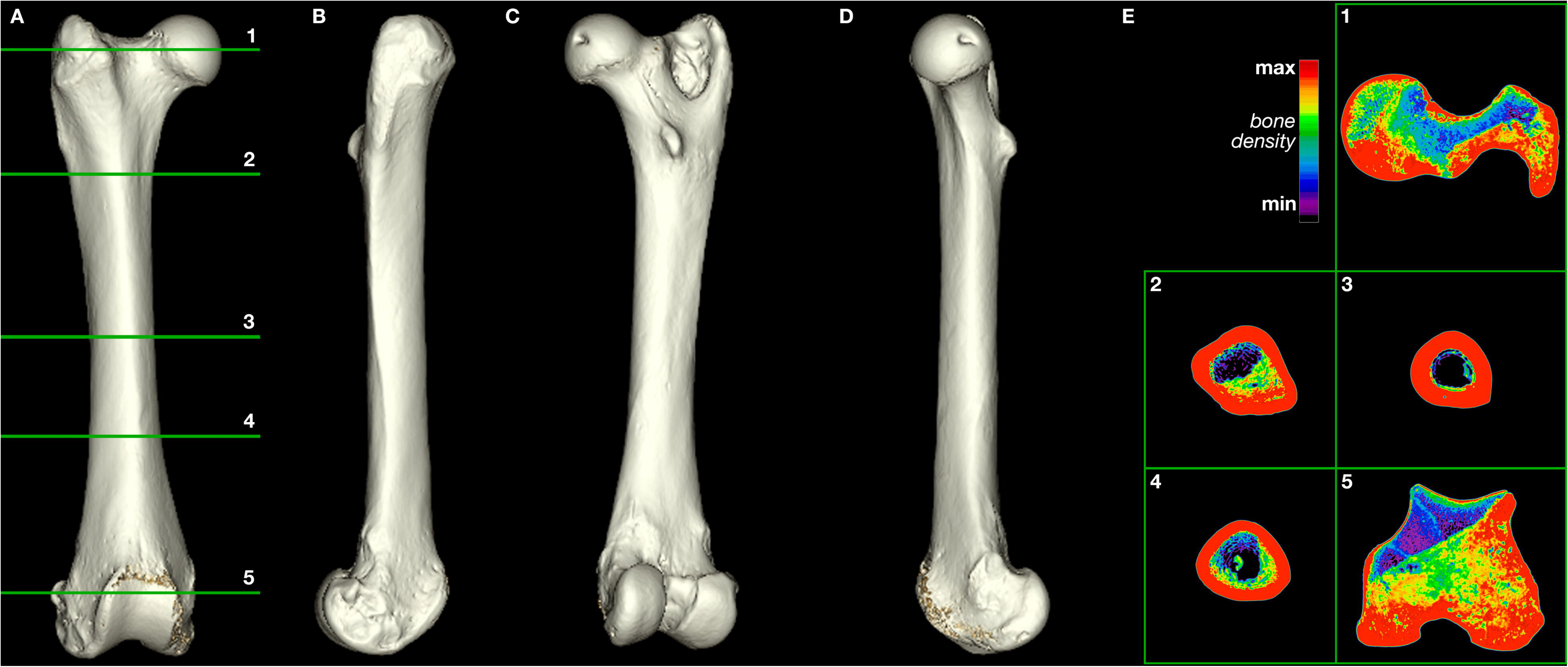
Metric data for the pathological *Smilodon* pelvis (LACMHC 131) in comparison to non-pathological specimens also from Pit 61/67 and potentially dysplastic pelvis specimens from other deposits. Inset diagram illustrates lengths measured. Across all measurements, the pathological pelvis lies well within the size range of non-pathological adult *Smilodon* pelves, suggesting that the animal to which it belonged also had grown to adult age. Numeric values are in Supplementary Data S4. **(A)** Bivariate plot showing ilium length (from the anterior-most point of the pelvis to the anterior border of the acetabulum) against total anteroposterior pelvis length. **(B)** Bivariate plot showing ischium length (from the posterior-most point of the pelvis to the posterior border of the acetabulum) against total anteroposterior pelvis length. **(C)** Bivariate plot showing ischium length against ilium length.

**Figure 7.**
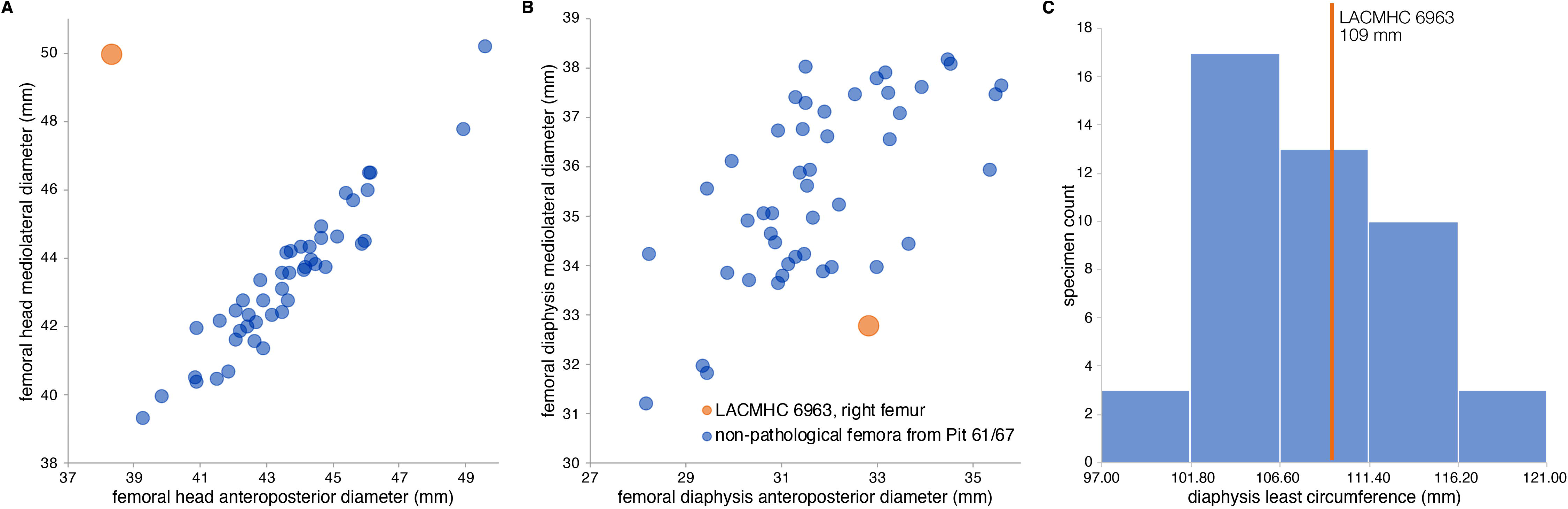
Metric data for the pathological *Smilodon* femur (LACMHC 6963) in comparison to non-pathological adult specimens also from Pit 61/67. Numeric values are in Supplementary Data S5. **(A)** Bivariate plot of mediolateral diameter against anteroposterior diameter of the femoral head. The femoral head of LACMHC 6963 is mediolaterally elongated and anteroposteriorly compressed compared to non-pathological specimens. **(B)** Bivariate plot of mediolateral diameter against anteroposterior diameter of the femoral diaphysis. The diaphysis of LACMHC 6963 is mediolaterally compressed relative to its anteroposterior diameter. **(C)** Histogram of femoral diaphysis least circumference, showing LACMHC 6963 (orange line) well within the range of non-pathological adult specimens and suggesting that the individual with the pathology also had reached adult age.

Using published regressions of body mass on postcranial measurements in living big cats^10–12^, we estimated the body mass of the individual bearing the pathologies. Our calculated results range from 114 kg (based on lengths on the right pathological innominate) to 125 kg (based on lengths on the left non-pathological innominate; likely an underestimate because big cats like the African lion comprise a minority of the broad comparative sample^10^), to 229 kg (based on femur circumference^11^) to 309 kg (using a different equation also based on femur circumference^12^). These estimates, like the skeletal measurements on which they are based, fall well within the range of body-mass estimates for non-pathological *Smilodon* from the same fossil deposit: from a median of 106 kg (median absolute deviation 14.15 kg) based on pelvis lengths^10^; to a median of 223 kg (median absolute deviation 24.95 kg) ranging up to 300 kg (median absolute deviation 36.12 kg) based on femur circumference^11,12^.

### Prevalence at Rancho La Brea

The focal specimens are from RLB deposit (“pit”) 61/67, which has produced 225 cataloged *Smilodon* pelvis specimens (99 left, 115 right, 13 complete; yielding a minimum number of individuals (MNI) = 128). Of these 225 pelves, 28 (14 left, 11 right, three complete; MNI = 17) bear lesions notable enough to have warranted inclusion in the Pathology collection of the museum at La Brea Tar Pits^9^. Three other Pit 61/67 pelves in the Pathology collection show deformations like those on the pelvis central to this study. These are: LACMHC 8904 (a left innominate; Fig. 13 in ^7^), LACMHC 8916 (a left innominate; Supplementary Fig. S3, this study), and LACMHC 9854 (a pelvis preserving both sides, described by Shermis^8^). Findings include osteoarthritic alteration of and chronic traction injuries around a shallow acetabulum, although etiology of these pathologies would need to be confirmed with CT imaging as in Figs. 4 and 5. Pit 61/67’s total of four potentially dysplastic pelves (MNI = 4) brings the prevalence in this deposit to a conservative estimate of 4 / 128 = 3.13%. All RLB deposits together catalog 1,624 *Smilodon* pelvis specimens (744 left, 772 right, 108 complete; MNI = 880); of these, we found three additional specimens with possible dysplasia (Supplementary Figs. S4-S6). This brings the dysplasia prevalence across all deposits to a conservative estimate of 7 / 880 = 0.80%. Other dysplastic specimens may have escaped inclusion in the Pathology collection, so this inferred percentage represents a minimum potential incidence of dysplasia among *Smilodon* individuals preserved at RLB.

Similarly, of 714 cataloged *Smilodon* femora (381 left, 333 right; MNI = 381) in Pit 61/67, six specimens (two left, four right) preserve skeletal pathologies notable enough to have warranted inclusion in the Pathology collection. However, LACMHC 6963—the pathological femur in the current study—is the only obviously dysplastic femur from Pit 61/67 in the Pathology collection. Indeed, we are not aware of femora in any other RLB deposit that have a femoral head as misshapen as that of LACMHC 6963. However, distortion likely would be less apparent in incipient stages of dysplasia, and such specimens may have escaped notice in the thousands of RLB specimens that have been curated and databased over the last century.

## Discussion

The arthritic degeneration visualized in the pathological *Smilodon* specimens could have arisen from one of three etiologies: traumatic, infective or degenerative arthritis. Findings on the specimens make infective or traumatic arthritis less likely. In the case of infective arthritis, the presupposition is that the animal developed typically before an insult that led to infection and subsequent obliteration of the hip joint. This assumption also holds true for the case of traumatic arthritis following an injury or fracture. However, the anatomical distortions of the right femoral head, in conjunction with the obliteration of the right acetabulum, suggest chronic changes that led to degeneration over time (Figs. 3, 4). The degeneration of the femoral head would not be expected if the degenerative change in the hip joint were due to infection or trauma, as the development of the pelvis and femur presumably would have been complete before the insult or injury occurred during the adult cat’s life.

Instead, the condition of the right acetabulum and right femoral head demonstrates anatomy consistent with developmental distortion. Typically, the head of the femur develops in conjunction with the acetabulum of the pelvis^13^. The spherical femoral head fits into the concentric-shaped acetabulum to form a ball-and-socket joint that enables a four-legged mammal to ambulate, lie down, sit down, stand up, and generally function normally^13^. In developmental hip dysplasia, however, the acetabulum does not develop appropriately, and the articulation between the femoral head and acetabulum is lost. An elliptical (as opposed to concentric-shaped) acetabulum causes progressive subluxation (dislocation) of the femoral head^14^, which can result in coxa plana, or necrosis of the bony nucleus of the femoral head^13^. This subsequent coxa plana produces flattening and degeneration of the normally spherical femoral head^15^.

Proper anatomical development and ossification of the hip joint rely on continuous and symmetrical pressure of the femoral head on the acetabulum, and dysplasia results from improper positioning of the femoral head within the acetabulum^13^. Dysplastic hips are characterized by pathological restructuring and accelerated remodeling of the joint in response to abnormal forces and tensions that create stress. This produces formation of new bone in some areas and resorption of bone in others, ultimately causing degenerative joint disease^13^. Dysplastic hips have varying degrees of deformity and malformation, but typically the acetabula are hypoplastic and deficient in various planes and dimensions (Supplementary Fig. S7).

This pathology starts to impact movement at the time of first walking, although minimal pain tends to ensue at this time because of the animal’s flexibility at its early age. As the joint cartilage wears out, however, bone begins to rub on bone. The resulting forces make the bone stiffer, producing osteophytes or bone spurs as well as sclerosis that manifests on CT imaging as increased bone density (Figs. 3 and 4; Supplementary Video S2; Supplementary Data S1-S2). At this point, loading the limb would cause pain, and range of motion would be limited. Therefore, the animal examined in this study would have spent as little time as possible on its right hindlimb, needing to compensate for the handicap by increasing the load on its left hindlimb. This compensation would explain the exostoses on the left ilium anterodorsal to the non-pathological acetabulum (Fig. 1; Supplementary File S1; Supplementary Video S1), indicating abnormal pulling of the quadriceps femoris muscles originating in this area.

Hip dysplasia is a heritable, polygenic condition that affects a range of mammal species^13^, including humans^14^. Canine hip dysplasia (CHD) is one of the most prevalent orthopedic diseases in domestic dogs (*Canis lupus familiaris*)^16^ and is very well studied, in part because it is similar to developmental dysplasia of the human hip^17^. Feline hip dysplasia (FHD) has received less clinical attention than CHD, possibly because its functional impairment is less overt or because domestic cats (*Felis catus*) are able to compensate for the resulting lameness better than dogs^18,19^. The overall results of physiological changes from dysplasia are mechanical imbalance and instability in the hip joint causing displacement due to opposing forces from the acetabulum and femoral head, and osteophytes in the acetabulum to compensate for cartilage loss^13^.

Embryologically, articular joints differentiate from skeletal mesenchyme *in situ* with the support of surrounding tissues that sustain mechanical and physiological forces that tend to pull on the joints^13^. In dogs, hip joints are normal at birth, as teratological factors and mechanical stresses that could displace the femoral head are rare at this time^13^. Epiphyseal ossification normally begins by 12 days of age; in dogs that eventually develop CHD, anatomical changes of the femoral head and pelvic socket begin before week three^20^. In dysplastic hips, the teres ligament, which is crucial for holding the femoral head in place, is too short; this produces luxation, or dislocation, of the top of the femoral head, beginning at around seven weeks^13^. This luxation increases throughout development, degrading the articular cartilage that surrounds the femoral head, delaying ossification of the femur and acetabulum^13^, and shortening the affected limb, as the femoral head becomes positioned higher in the acetabulum.

In clinical reports of hip dysplasia in domestic cats, osteoarthritis (i.e., degenerative joint disease, DJD) of the hip secondary to FHD is well known^21^. For example, osteoarthritis was recorded in 43 of 45 (95.6%) of cats with FHD^22^. As well, in 5 of 13 (38.5%) cases of hip osteoarthritis with a radiographically or historically identifiable cause, hip dysplasia was pinpointed as the cause, with the remaining cases resulting from trauma or equivocal between trauma and dysplasia^18^. A recent study of FHD in Maine Coons—a large-bodied domestic cat breed in which hip dysplasia is known to be common—calculated a prevalence of 37.4%, finding severity to increase with age and body mass^23^. The same study further highlighted a genetic correlation between FHD and large body size within the Maine Coon^23^, inviting inquiry into how FHD impacts other breeds and non-domestic felid species across a range of body sizes.

Reports of FHD in non-domestic large cats are rarer than in domestic cats. Captive snow leopards have exhibited hip dysplasia; across 14 zoos, seven cases were classified as moderate to severe, and at least two individual snow leopards needed total hip replacement before being able to breed^24,25^. Accounts of hip functional impairment in other captive large cats have tended to report osteoarthritis, which can be associated with FHD though may also stem from trauma and increased age^26–28^.

For wild-caught large cats, the only comprehensive study of which we are aware is a survey of 386 individuals (283 wild-caught) across three felid genera mounted as exhibit skeletons in multiple North American natural history museums^27^. Though not focusing on hip dysplasia, the study tracked degenerative joint disease, which may be associated with dysplasia^18,22^. The sample recorded DJD in 9.7% of 31 tigers, 2.3% of 88 African lions, and 5.1% of 59 mountain lions (*Puma concolor*), and none in five other species of big cat. These frequencies are low compared to domestic cats, perhaps owing to differences in body size, diet, and lifestyle between large wild cats and domestic cats, as well as selective breeding constraining genetic variation in domestic animals. Furthermore, selection against hip dysplasia would be expected in the wild because hip dysplasia would compromise hunting^21^. Though this study identified instances of non-inflammatory osteoarthritis in the shoulder, elbow, and stifle joint, it found none in the hip. However, 4% of all joints afflicted by spondyloarthropathy—a form of inflammatory arthritis—included the hip^27^.

*What is the significance of Smilodon, an extinct Pleistocene predator, having the same congenital defect as living domestic cats and dogs?* Previous workers have inferred social behavior from *Smilodon’s* pathologies, interpreting signs of healing as evidence that the animal continued to live after injury^9^. Given the severity of many injuries, authors have argued, the animal would have starved to death had it not operated within a social structure. The present hip dysplasia having manifested from a young age—hindering the animal’s ability to hunt prey and defend a home range over the course of its life—supports this assertion, although other inferences are possible.

Sociality, the degree to which individuals live with conspecifics in groups^29^, is difficult to infer in *Smilodon* given that it has no living analogues or closely related taxa. Estimated to have weighed between 160 and 350 kg (^3,11,^ this study), *Smilodon* was at least the size of the Amur tiger (*Panthera tigris altaica*), the largest living cat; some estimates reach 369 to 469 kg, placing *Smilodon* in the range of the largest extant ursids^12,30^. No living felid has *Smilodon*’s elongate, knife-like canines or stocky, powerful build. As well, *Smilodon* (of the extinct felid lineage Machairodontinae) is only distantly related to extant large felids (Felinae), introducing further uncertainty. Based on its robust morphology (e.g., ^31,32^) and on evidence from stable isotopes (e.g., ^4^), it likely stalked and ambushed prey; therefore, it may have been comparable to the African lion (*Panthera leo*), which has a similar hunting strategy and is the only truly social extant felid^33^. Yet sociality varies across felid species, including within a genus; for example, other extant pantherines like tigers (*P. tigris*) show incipient sociality^34^, while jaguars (*P. onca*) are solitary except for females with cubs. Social strategies also can vary within species, e.g., between sexes. For instance, African lion females are philopatric and social throughout their lives, while adult males are often nomadic and solitary until joining a gregarious pride, which itself usually lasts for only a few years^35^. This social variation complicates behavioral inferences based on ancestral reconstructions.

Advocates of the solitary-cat hypothesis^36,37^ have cited *Smilodon*’s small relative brain size determined using endocranial casts as support for solitary behavior, because sociality exerts high cognitive demands. However, in 39 species across nine carnivoran families, larger relative brain size was found to correlate with problem-solving capabilities rather than social behavior^38^. Rather than analyses of overall encephalization across carnivoran families, studies of relative regional brain volume within families and species have been more informative regarding sociality^39,40^. In both African lions and cougars (*Puma concolor*, a solitary species), total relative endocranial volume was not sexually dimorphic; however, relative anterior cerebrum volume was significantly greater in female African lions than males, a difference absent in cougars^35^.

Though regional endocranial studies have yet to be performed on *Smilodon*, the gregarious-cat hypothesis has drawn support from multiple lines of evidence. One is the abundance of *Smilodon* relative to prey at RLB^7,8,31^, although detractors have pointed out that some extant large cats aggregate at carcasses despite otherwise being solitary^37^. A full range of ages is present among RLB *Smilodon*; in contrast, animals interpreted to be solitary, such as the American lion *Panthera atrox,* are represented largely by adult individuals^41^. As well, the proportions of social and solitary species at RLB parallel those drawn to audio recordings of herbivore distress calls in the African savanna, suggesting that RLB *Smilodon* sample sizes are more consistent with it having been social rather than solitary^42,43^. The lack of size sexual dimorphism in *Smilodon* is more typical of modern solitary cats^44^ but could also be reflective of monogamy within a gregarious species, like modern wolves. Most relevant to the current study, the existence of healed injuries in *Smilodon* also has been interpreted as evidence for social behavior, with the assumption that surviving long after serious injury would be difficult if not impossible without cooperative sociality^9^. We now revisit this interpretation considering the novel diagnosis of hip dysplasia in this study.

*Smilodon*’s large body size necessitated preying on megaherbivores for adequate sustenance^3^. To do so, like most large cats today, it would have used its hindlimbs for propulsion and acceleration^45,46^, a pounce behavior enabled by its morphology*. Smilodon’s* ratio of total forelimb to hindlimb length is greater while its ratio of tibia to femur length ranks lower than in living felids^31^. The shorter hindlimbs lacking the distal limb elongation in cursorial animals suggest that *Smilodon* was an ambush predator surpassing the ability of felids today^47^. Hunting large prey is dangerous^48^. After the initial hindlimb-powered leap, *Smilodon* would have grappled with its struggling prey, as evidenced by traumatic injuries in the rotator cuff and radiating from the ventral midline dorsolaterally to where the ribs articulate with the spine^5^. As it subdued prey with robust forelimbs^32,45^ under enough torque to injure the lumbar vertebrae^5^, *Smilodon* would have needed to leverage itself against the ground using its hindlimbs. Therefore, the pelvis and femur would have been critical to multiple phases of its hunting strategy.

A dysplastic individual would have encountered much difficulty hunting in this manner. Yet, as evidenced by the complete fusion of its pelvic and femoral epiphyses (Figs. 1, 2) as well as its large body size (Figs. 6, 7), the individual in this study had reached adult age. (Studies of the detailed timing of epiphyseal fusion in large wild cats are lacking, but distal femoral epiphyses fuse at around the same time as or soon after proximal femoral epiphyses in domestic cats and dogs^49,50^. Given this, the broken distal femur likely had a fused epiphysis, as on its intact proximal end.) Limbs in African lions completely fuse between 4.5 and 5.5 years^51–53^, so it is reasonable to assume that adulthood in *Smilodon* likely started at around four years old. This estimate is reinforced by bone histological work quantifying at least four to seven lines of arrested growth (LAGs; one per growth year) in limb bones with fused epiphyses belonging to *Smilodon fatalis* from the Talara asphaltic deposits in Peru^54^. Some LAGs in the Talara histological specimens likely have been masked by secondary bone remodeling, which may be more extensive in larger-bodied taxa^54^, making these specimens possibly older than the number of visible LAGs suggest. Therefore, four years represents a likely minimum age for this individual, although it could have been much older.

Ontogenetic growth patterns in teeth and bone further support inferences of sociality. In *Smilodon*, teeth appear to mature earlier than when sutures and long-bone epiphyses fuse, suggesting delayed weaning, prolonged juvenile dependence, and extended familial care until the adult hunting morphology—saber canines and robust limbs—was complete^44^. At RLB, most sampled *Smilodon* specimens show significant pulp cavity closure of the lower canine (14 of 19 specimens over approximately 80% closure), a sign of dental maturation^55^. This contrasts with RLB pantherine pulp cavities, which are more evenly distributed across the closure percentage range, suggesting that teeth mature earlier in *Smilodon* than in pantherines. (Other age assessments have ruled out the possibility that *Smilodon* juveniles were underrepresented relative to pantherines^41^.) At Talara, age determination by dentition yields low estimates of juveniles (zero based on skulls; 8% based on dentaries), but age determination based on limb epiphyseal fusion yields higher estimates (41% juveniles)^54^. Histology of Talara *Smilodon* long bones reinforces this mismatch, as an apparent adult femur with fused epiphyses and seven LAGs was found to lack avascular and acellular subperiosteal lamellar bone^54^, suggesting that it had not yet finished growing. Further, prolonged parental care was interpreted in a recent description, from Pleistocene deposits in Corralito, Ecuador, of two subadult *Smilodon fatalis* individuals inferred to have been siblings and associated with an adult that was likely their mother^56^. This scenario of prolonged parental care, like that in the social African lion, would help explain how the individual in this current study survived to adulthood given its debilitating handicap.

Novel application of CT visualization to an old question of paleopathology has enabled diagnosis of hip dysplasia, a lifelong condition, in an individual *Smilodon fatalis* saber-toothed cat. This individual was likely not the only *Smilodon* afflicted with hip dysplasia: multiple RLB *Smilodon* pelvic specimens, especially that described by Shermis^8^, exhibit gross morphology similar to the pathological pelvis examined in this study (Supplementary Figs. S3-S6). The individual examined in this study reached adulthood (at least four to seven years of age) but could never have hunted nor defended territory on its own, given its locomotor impairment that would have been present since infancy. As such, this individual likely survived to adulthood by association with a social group that assisted it with feeding and protection.

Further conclusions are limited by the lack of a comprehensive and systematic comparative dataset comprising pathological postcrania from extant species, a persistent limitation of paleopathological studies^5^. Natural history museums may acquire cranial remains from zoos or similar institutions but often lack storage to accommodate postcranial skeletons, especially for large mammals. As well, while radiographic studies on domestic cats and dogs illustrate the nature of hip dysplasia, these studies tend to examine pathological bones *in situ*, still embedded in a muscular framework (e.g., Supplementary Fig. S7). This is opposed to the bones-only, flesh-free context of paleopathological specimens. Computed tomography and digital data may be key to building a comparative paleopathology dataset in the future.

Within the scope of this study, we cannot rule out the hypothesis that the pathological animal was a scavenger and may have obtained food outside the context of a social structure. It is also possible that, regardless of its disability, its large size and fearsome canines made it a strong interference competitor. However, the pathological specimens examined here are consistent with the predominance of studies supporting a spectrum of social strategies in this extinct predator. In many extant carnivorans, sociality offers the benefits of cooperative hunting and rearing of young (e.g., ^57^): benefits that likely also applied to *Smilodon* in the late Pleistocene. As *Smilodon* coexisted with a rich megafaunal carnivore community including dire wolves (*Aenocyon dirus*), American lions (*Panthera atrox*), and short-faced bears (*Arctodus simus*), cooperative sociality may have aided its success as a predator in a crowded field.

## Methods

All specimens examined are from the Pleistocene-age Rancho La Brea collections of the museum at the La Brea Tar Pits (LBTPM), part of the Natural History Museums of Los Angeles County (NHMLAC; formerly LACM), Los Angeles, California. At Rancho La Brea, different fossiliferous asphaltic deposits (which became human-made “pits” during the historical excavation process) resulted from periodic asphalt seep activity, entrapping organisms across different timespans over the past 55,000 years^58^ with varied depositional environments and taphonomic histories^59^. In this context, we reduced potential variability in these factors by selecting all specimens from a single deposit, Pit 61/67. Pit 61/67 is the most recent deposit at Rancho La Brea that precedes the late Pleistocene megafaunal extinctions at around 11,000 years before present^60^, at which point *Smilodon fatalis* became extinct.

We examined the external surfaces of the pathological pelvis including the right innominate (LACMHC 131) and associated pathological right femur (LACMHC 6963). We also inspected in detail an unassociated non-pathological right femur (LACMHC K-3232) from the same deposit and of similar size and ontogenetic stage. Initial surface scanning of all specimens was carried out at LBTPM using an Artec Space Spider (Artec 3D) as a means of digital preservation and to provide a 3D visual with color. The surface scans were processed in Artec Studio 12 and fused into a model with a resolution of 0.2 mm. CT imaging of the three specimens was performed at the S. Mark Taper Foundation Imaging Center, Los Angeles, California, on a GE Revolution (GE Healthcare, Waukesha, WI) 256-slice scanner with 0.625 mm slice thickness. Imaging parameters were KVP=120, mA=300, 0.5 second rotation time, and 0.51 pitch using a medium body FOV. The data were acquired in the axial plane, reformatted into soft tissue and bone algorithms, and viewed in the axial, coronal, and sagittal planes. CT images were converted to 3D models using the segmentation software Mimics (Materialise). Geomagic Freeform (3D Systems) was used to upload and determine placement of the plane for cross-sections of the 3D reconstructions.

To estimate the individual’s body size and the population incidence of this pathology, we measured pelvis lengths and femur circumference on the pathological specimens and compared them to non-pathological pelves and femora from Pit 61/67. Lengths were measured using Mitutoyo calipers. We restricted this comparison to Pit 61/67, source of the pathological specimens, because previous work has revealed body-size differences among *Smilodon* from different RLB deposits, likely due to climate adaptation^61^. We estimated body size using published regressions of pelvis lengths^10^ and femur circumference^11,12^ against body masses of living felids like the African lion.

While quantitative methods for scoring hip dysplasia exist (e.g., ^62^), these methods were developed for veterinary use on live animals presumably smaller than Pleistocene megafauna, employed to describe x-rays of articulated limbs (e.g., ^23^). Therefore, this scoring system was inappropriate for the disarticulated and exposed bony material in the paleontological record, as in this study. However, several RLB *Smilodon* pelvic specimens preserve gross morphology consistent with dysplasia, which we surveyed to calculate prevalence of this pathology in the RLB *Smilodon* population.

No permits were required for the described study.

## Acknowledgements

We acknowledge that Rancho La Brea is on the traditional territory and homelands of the Chumash and Gabrielino / Tongva people. We thank Aisling Farrell at LBTPM for facilitating specimen access and loans; CT segmentation specialists at the S. Mark Taper Foundation Imaging Center for CT-scanning specimens; Jeff Busey and Jessica Butler at Zimmer Biomet for aiding design of 3D models; Karin Rice for x-ray images of Erik the cat; Fred Heald for his work on the LBTPM Pathology collection; and Andrew Kitchener, Carlo Meloro, Alexis Mychajliw, Blaire Van Valkenburgh, and an anonymous reviewer for constructive comments. A U.S. National Science Foundation Postdoctoral Research Fellowship in Biology (DBI-1812301) funded MAB.

## Author Contributions

MAB and AKS wrote the manuscript text; CMH surface-scanned specimens and created videos; MAB, CMH, and AKS prepared figures; MAB collected quantitative data; CAS described comparative-specimen pathologies; RK and ELL acquired CT scans; MAB and ELL guided study design. All authors reviewed and approved the manuscript.

