## Supplementary Information for "Computed tomography reveals hip dysplasia in the extinct Pleistocene saber-tooth cat *Smilodon*"

Supplementary File (on Dryad: <https://doi.org/10.6071/M3S09M>)

Supplementary Videos (on Dryad: <https://doi.org/10.6071/M3S09M>)

Supplementary Data (on Dryad: <https://doi.org/10.6071/M3S09M>)

Supplementary Figures (in this document)

**Supplementary File S1.** Three-dimensional PDF of the pathological pelvis (LACMHC 131) and femur (LACMHC 6963).

**Supplementary Video S1.** A video movie of a structured light surface scan of LACMHC 131, the pathological pelvis belonging to *Smilodon fatalis*. Movie created in Adobe Photoshop CC by Carrie Howard.

**Supplementary Video S2.** A video scrolling anteroposteriorly through LACMHC 131, the pathological pelvis belonging to *Smilodon fatalis*. Ventral is at top. Video created by Carrie Howard from CT scans generated at the S. Mark Taper Foundation Imaging Center at Cedars Sinai Medical Group.

**Supplementary Data S1.** Compressed zip file containing the full computed tomography scan of LACMHC 131, the pathological pelvis.

**Supplementary Data S2.** Compressed zip file containing the full computed tomography scan of LACMHC 6963, the pathological femur.

**Supplementary Data S3.** Compressed zip file containing the full computed tomography scan of LACMHC K-3232, the non-pathological right femur.

**Supplementary Data S4.** CSV file containing basic morphometric data for comparative *Smilodon fatalis* pelvis specimens, both pathological and non-pathological. LACM = Los Angeles County [Natural History] Museum; HC = Hancock Collection.

**Supplementary Data S5.** CSV file containing basic morphometric data for comparative *Smilodon fatalis* femur specimens, all non-pathological. LACM = Los Angeles County [Natural History] Museum; HC = Hancock Collection.

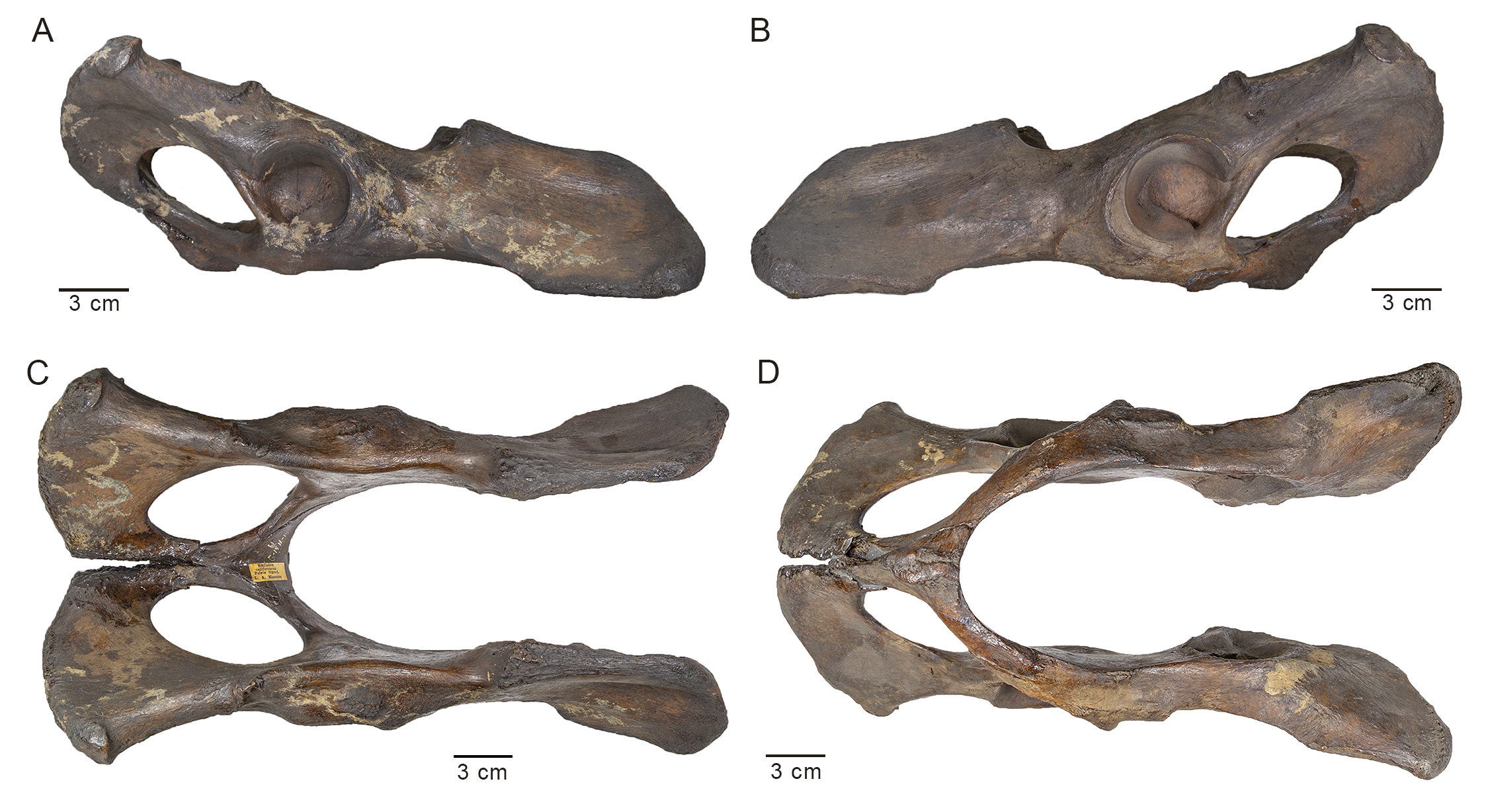

**Supplementary Fig. S1.** Photographs of LACMHC K-2584, a non-pathological pelvis belonging to *Smilodon fatalis*. **(A)** Lateral view of right side; anterodorsal end to the right. **(B)** Lateral view of left side; anterodorsal end to the left. **(C)** Dorsal and **(D)** ventral views; anterior end to the right.

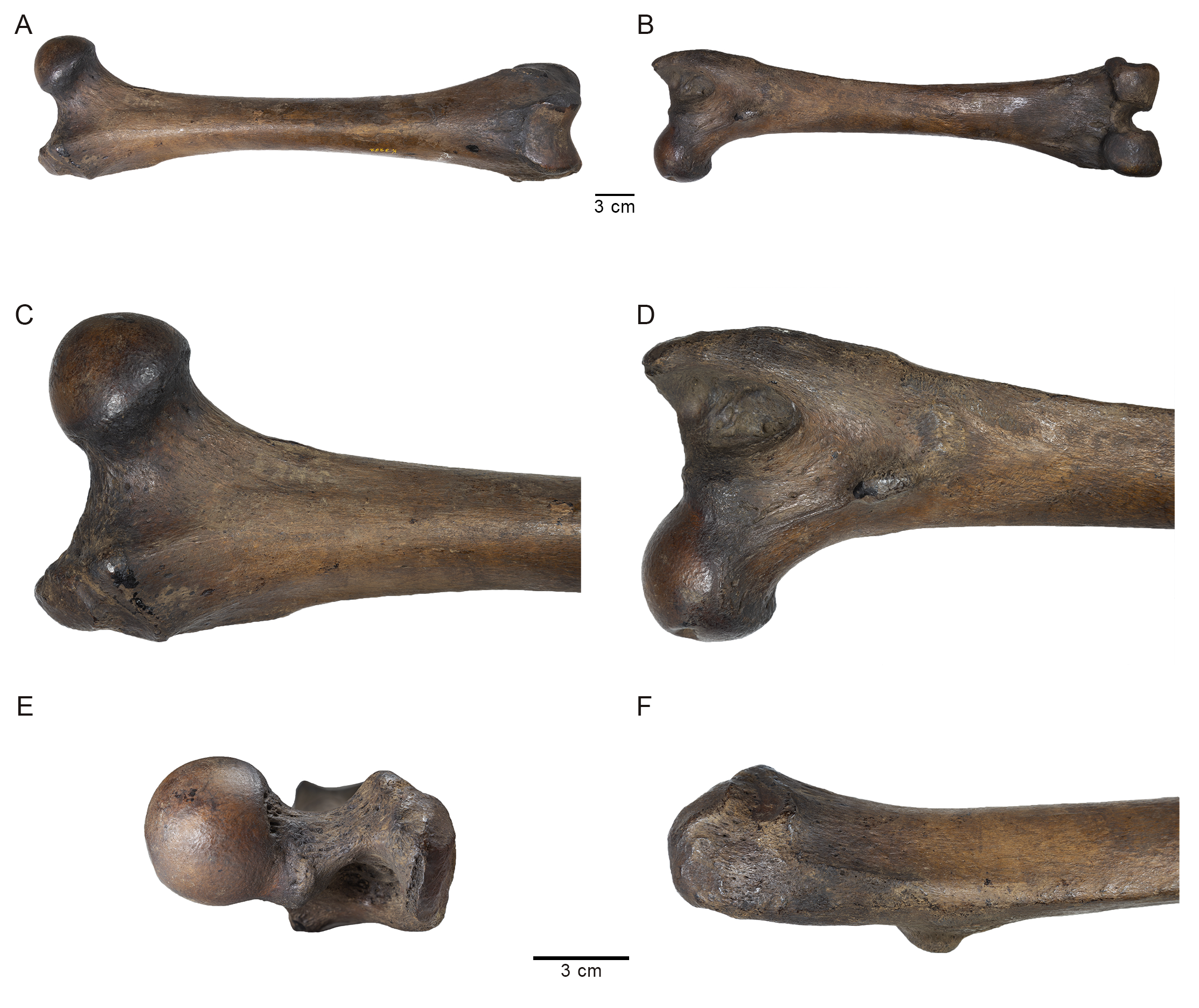

**Supplementary Fig. S2.** Photographs of LACMHC K-3232, a non-pathological right femur belonging to *Smilodon fatalis*. **(A)** Anterior and **(B)** posterior views of full femur; proximal end on the left. **(C)** Anterior and **(D)** posterior close-up views of the proximal end, including the spherical femoral head, greater trochanter, and lesser trochanter. **(E)** Dorsal close-up view of the femoral head, greater trochanter, and lesser trochanter in lower center background. **(F)** Lateral close-up view of the greater trochanter and lesser trochanter (lower center). The upper scale bar refers to A and B, and the lower scale bar refers to C, D, E, and F.

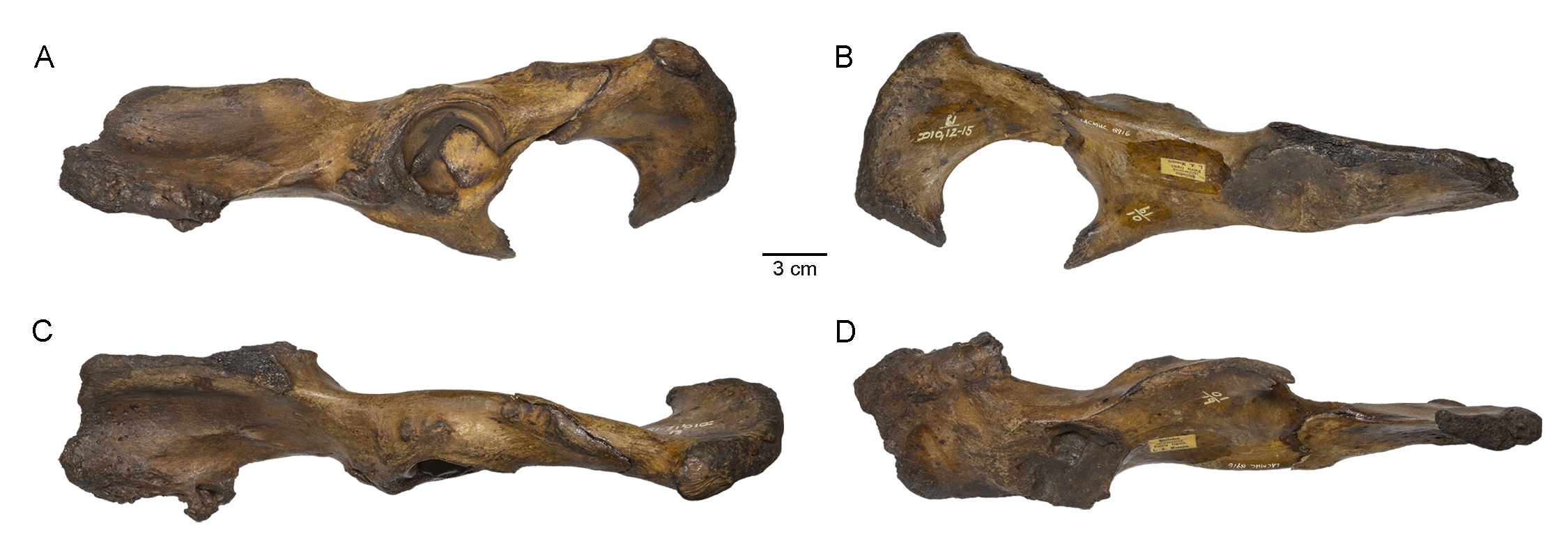

**Supplementary Fig. S3.** Photographs of LACMHC 8916, a pathological left innominate of *Smilodon fatalis* from Pit 61/67 exhibiting potential signs of dysplasia, in particular heavy exostoses and possible traction injuries in the origins of the *M. rectus femoris* and *M. iliacus*. **(A)** Lateral view; anterodorsal end to the left. **(B)** Medial view; anterodorsal end to the right. **(C)** Dorsal and **(D)** ventral views; anterior end to the right.

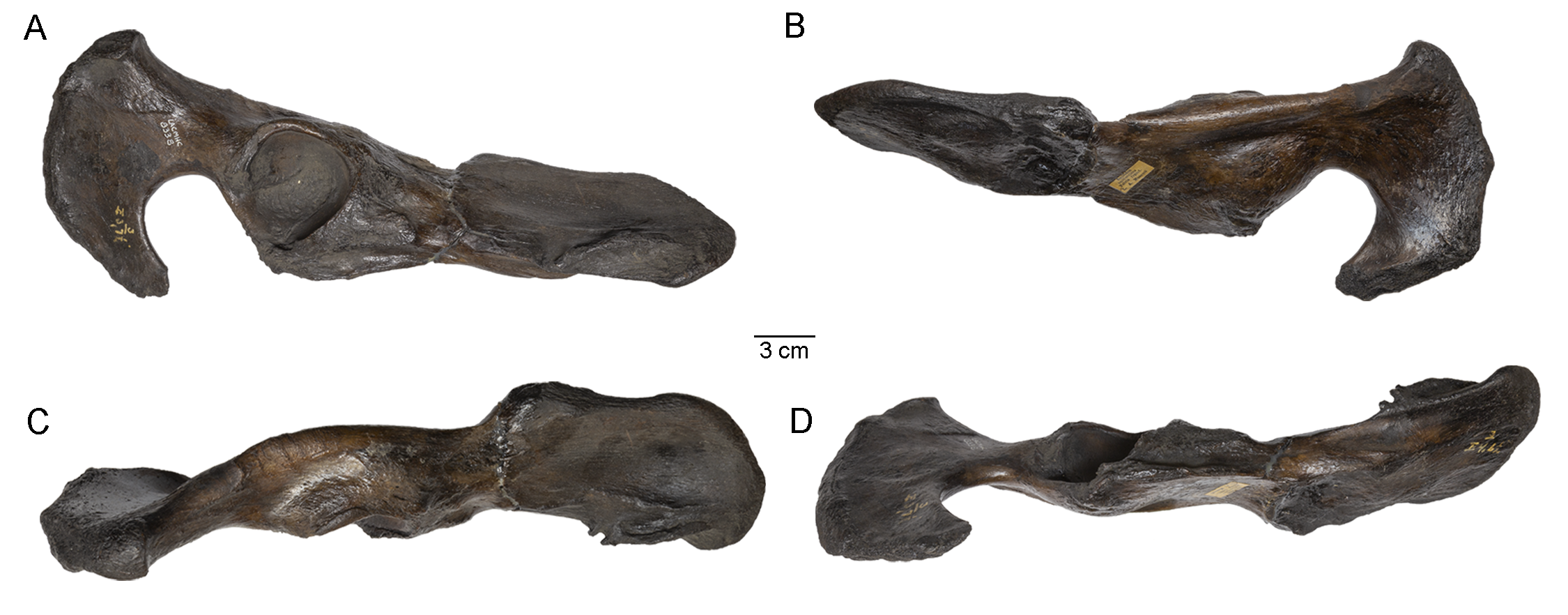

**Supplementary Fig. S4.** Photographs of LACMHC 8338, a pathological right innominate of *Smilodon fatalis* from Pit 3 exhibiting potential signs of dysplasia. Deepening of the acetabulum has caused a bulge in the medial wall, likely impacting gait and weight-bearing, and the origins of the *M. iliacus* and *M. gluteus medius* exhibit traction injuries. **(A)** Lateral view; anterodorsal end to the right. **(B)** Medial view; anterodorsal end to the left. **(C)** Dorsal and **(D)** ventral views; anterior end to the right.

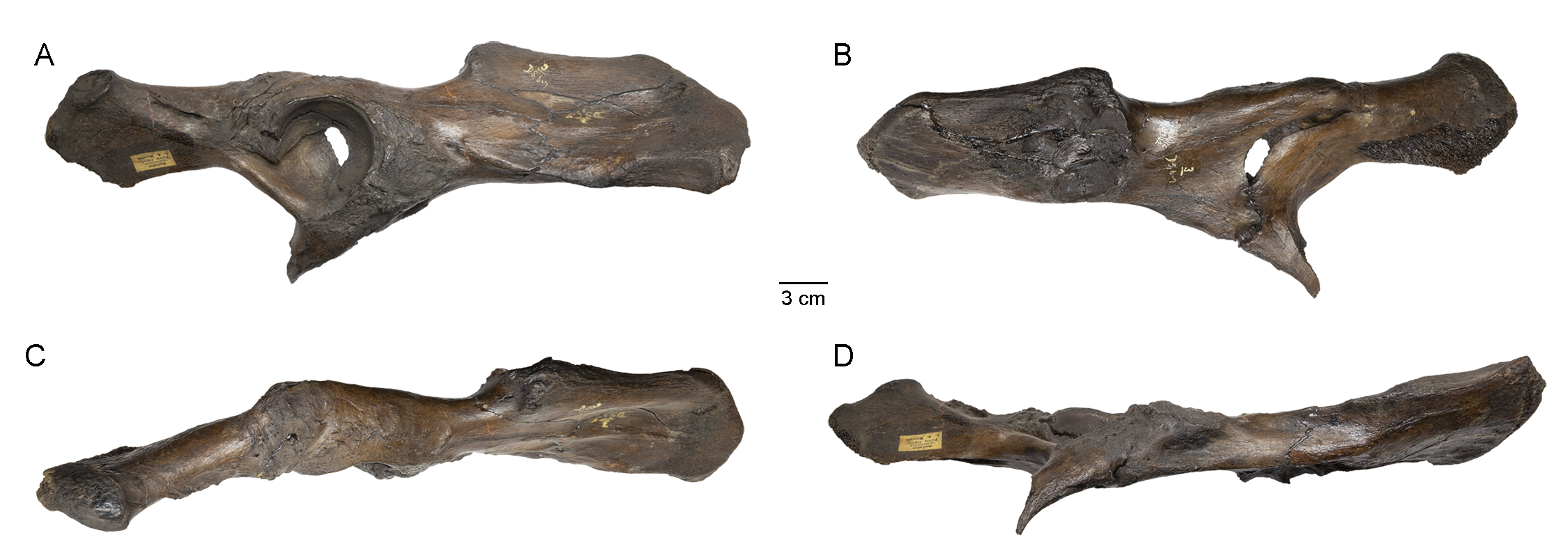

**Supplementary Fig. S5.** Photographs of LACMHC 8342, a pathological right innominate of *Smilodon fatalis* from Pit 3 exhibiting potential signs of dysplasia. A healed fracture crosses the acetabulum; the associated callus build-up has united in a line involving both the pubis and the ischium at an oblique angle. The acetabular joint is intact and smooth, however, and the matching femur would be of great interest if found. **(A)** Lateral view; anterodorsal end to the right. **(B)** Medial view; anterodorsal end to the left. **(C)** Dorsal and **(D)** ventral views; anterior end to the right.

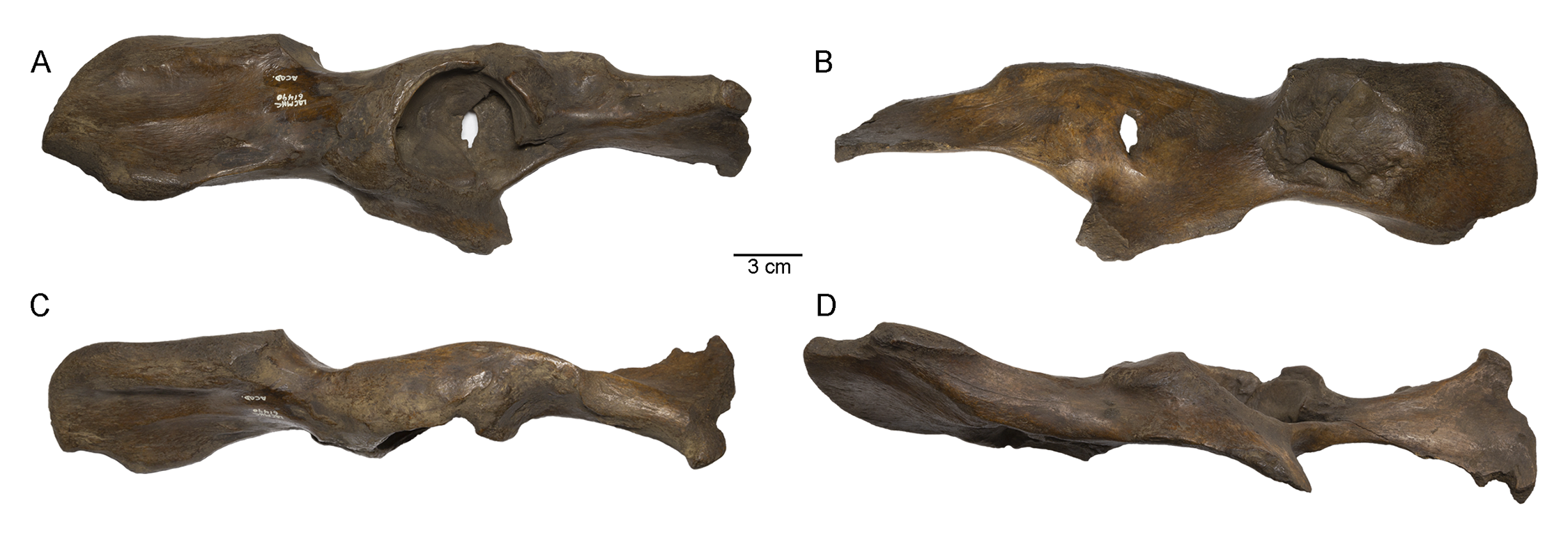

**Supplementary Fig. S6.** Photographs of LACMHC 61490, a pathological left innominate of *Smilodon fatalis* from Pit “Academy” exhibiting potential signs of dysplasia. The posterior wall of the acetabulum is wrinkled deeply, and the anterodorsal aspect exhibits an apparent incompletely healed fracture. CT imaging of the internal bone structure would aid in identifying etiology. **(A)** Lateral view; anterodorsal end to the left. **(B)** Medial view; anterodorsal end to the right. **(C)** Dorsal and **(D)** ventral views; anterior end to the right.

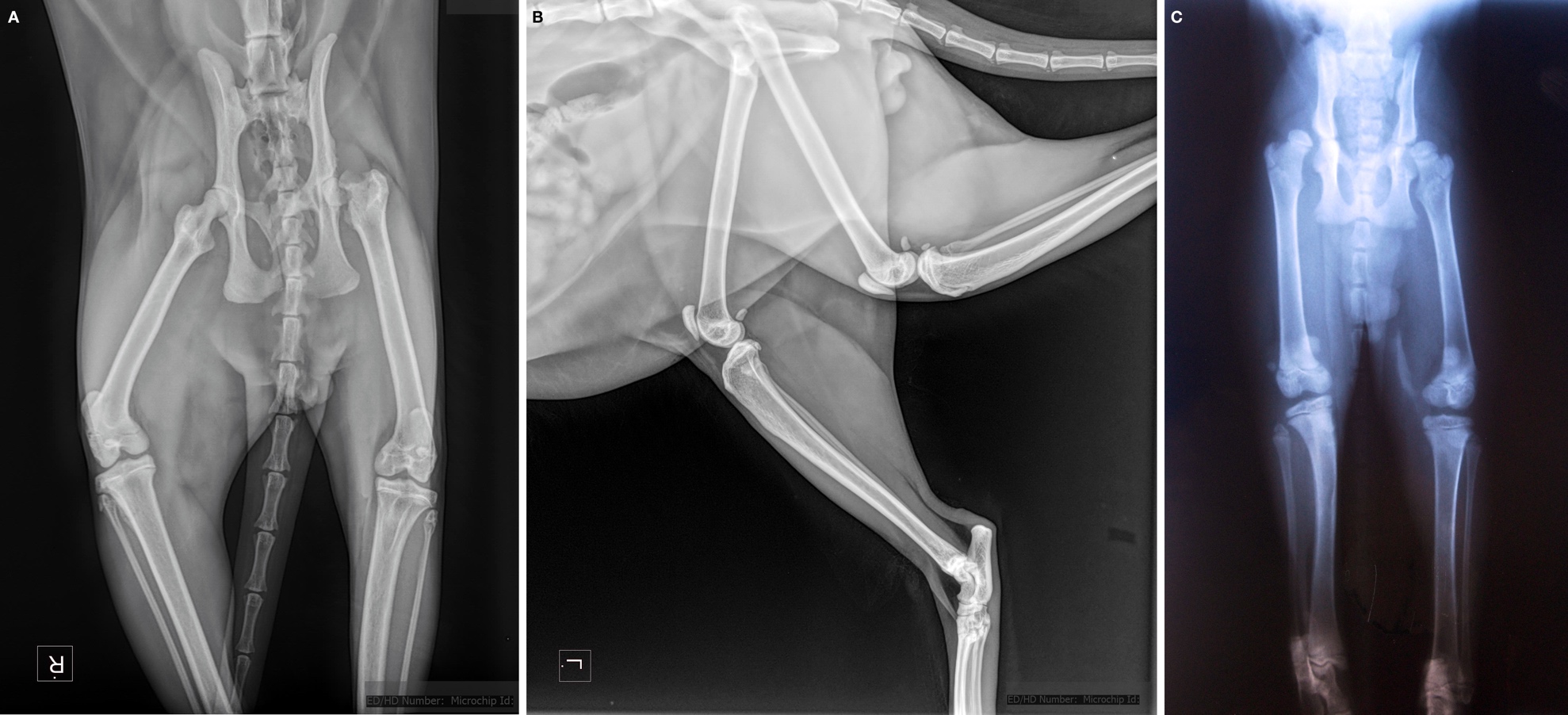

**Supplementary Fig. S7.** X-ray images of two domestic cats, both of which exhibit the radiographic hallmarks of hip dysplasia. (a) Ventrodorsal and (b) lateral radiographs of the pelvis and hip joints of an individual demonstrating severe left hip dysplasia. The hypoplastic and shallow left acetabulum, characteristic of dysplasia, has resulted in significant femoral head deformity, subluxation, and dislocation of the hip joint. Relative to the right hip, there is evidence of chronic remodeling and degenerative arthritic changes in the left acetabulum and femoral head, as well as shortening of the left leg and asymmetry between the two extremities. Additionally, the left femoral head is flattened and severely anatomically distorted. The lateral view demonstrates asymmetry in the left and right hip joints due to the dorsally dislocated left hip. (c) Ventrodorsal view of the pelvis and hip joints of another individual demonstrating signs of right hip dysplasia. The shallow right acetabulum does not provide adequate coverage for the right femoral head to fit concentrically, resulting in hip instability. The right femoral head is proximally dislocated, creating a high-riding hip. (a-b) courtesy of Carrie M. Howard, (c) courtesy of Karin A. Rice.
